# Sex-Dependent Alterations in Fusiform Face Area Activation During Emotional Face Processing in Autism

**DOI:** 10.64898/2026.05.27.728171

**Authors:** Sneha Lakamsani, Jeffrey Eilbott, Stefen Beeler-Duden, Kevin Pelphrey

## Abstract

Atypical processing of emotional faces has been proposed as a characteristic of Autism Spectrum Disorder (ASD), but functional neuroimaging research has yielded inconsistent findings. Prior work is limited in generalizability due to methodological heterogeneity, imbalanced or small sample sizes, and underrepresentation of females. The present study examined functional brain activation during the Hariri Emotional Face-Matching Task (EFMT) in a large, sex-balanced sample of both typically developing and ASD participants (n=295, 8-18 years old) from the multi-site Autism Center of Excellence GENDAAR project. Using an ROI-driven approach, we targeted the right FFA, right OFA, right pSTS, and bilateral amygdala, we investigated whether ASD diagnosis was associated with atypical regional activation when viewing emotional faces, and if these differences were generalizable across sexes. Results revealed a group-by-sex interaction in the right FFA, driven by divergence of ASD males from the ASD female and typically developing participants. Generally, ASD females did not diverge greatly from typically developing populations. These findings suggest that atypical face processing is present, but meaningfully modulated by sex, underscoring the importance of sex-balanced, well-powered developmental samples in autism.

## Introduction

Autism spectrum disorder (ASD) is a neurodevelopmental condition marked by differences in social communication and interaction, along with restricted or repetitive patterns of behavior and interests^1^. Because successful social interaction depends heavily on the rapid interpretation of faces, face perception has been a central focus of autism research for more than two decades^2^. Faces convey rich social information, including identity, gaze direction, affective state, and communicative intent, and efficiently extracting this information is fundamental to adaptive social functioning. Atypical engagement with faces has therefore been proposed as a mechanism contributing to the social phenotype of autism^3–5^.

Face processing emerges early in development and is one of the most robust specializations of the social brain^6,7^. Human infants preferentially orient to faces from the earliest days of life, and these early perceptual biases scaffold the development of increasingly sophisticated abilities to decode identity, emotion, and intention throughout childhood and adolescence. In typically developing individuals, these processes are supported by a distributed neural system with right hemisphere dominance^8–10^. The core system includes the occipital face area (OFA), fusiform face area (FFA), and posterior superior temporal sulcus (pSTS), as well as regions involved in salience detection and affective evaluation, such as the amygdala^11–13^. The right OFA is believed to aid in the initial structural encoding of facial features, which serves as input to regions further downstream^14,15^. The right FFA, preferentially responds to faces over other objects and supports identity recognition^8,12,13^. The right pSTS is responsible for processing dynamic aspects of faces—gaze direction, mouth movements, and emotional expressions^16,11^. The right-lateralization of these regions reflects the right hemisphere dominance of the face processing pathway, evidenced by converging evidence from prosopagnosia following right-hemisphere damage, split-brain studies, and functional neuroimaging^17,18^. In contrast, the extended face processing network recruits the amygdala bilaterally for affective processing and relevance detection, demonstrating the amygdala’s broader role in detecting emotional stimuli regardless of visual field^19,20^. Both the left and right amygdala are responsive to emotional faces, with the right amygdala potentially responding more to fearful faces while the left amygdala is preferential to prolonged emotional processing^21,22^. This network supports both the processing of invariant facial properties and the interpretation of dynamic, socially meaningful signals, such as emotional expressions and gaze.

Given the centrality of social-communication differences in ASD, this neural face-processing system has been a major focus of functional neuroimaging research. Despite more than two decades of study, the literature remains notably inconsistent. Some studies report reduced activation in key regions such as the FFA, pSTS, and amygdala, findings often interpreted as evidence of a core deficit in social perception^23–26^. In contrast, other studies find attenuated or minimal group differences when attention, gaze behavior, or task demands are considered^27–30^, while still others suggest that autistic individuals may recruit alternative or compensatory neural pathways during face processing^31,32,2,33^. These discrepancies may reflect differences in paradigm demands, sample composition, and analytic approach as much as any single underlying brain mechanism^2,34,35^.

A major contributor to this lack of consensus is methodological heterogeneity. “Face processing” encompasses a wide range of tasks that differ substantially in their cognitive and perceptual demands, including passive viewing, identity discrimination, gaze processing, and emotion recognition. These distinctions are critical because different tasks engage partially distinct components of the broader face and social-cognitive network^11,36^. In particular, paradigms that require explicit evaluation of emotional expressions are likely to recruit not only ventral visual regions but also pSTS, amygdala, and frontal systems involved in salience detection and socioemotional interpretation^25,37^. Consequently, variability in task design may obscure or exaggerate diagnostic group differences, limiting the interpretability and generalizability of prior findings.

A second limitation of the literature concerns sample composition. A series of seminal early fMRI studies laid the foundation for the modern neuroimaging literature on face processing in autism and were instrumental in identifying the FFA, pSTS, amygdala, and related socioaffective circuitry as candidate systems of interest^23–25,38–42^. These studies were highly generative and established many of the core questions that continue to shape the field. At the same time, as with much early neuroimaging work, they were often conducted in relatively modest sample sizes, limiting statistical power and precision in characterizing heterogeneity or developmental variation. This issue is especially consequential in developmental neuroimaging, where neural responses to social stimuli change across childhood and adolescence. Yet relatively few studies have examined large, developmentally informative samples spanning this critical period, leaving unresolved whether reported group differences generalize across development.

A third and particularly important limitation is the historical underrepresentation of females^43^. Although ASD is diagnosed more often in boys than in girls, growing evidence suggests that this discrepancy may reflect differences in ascertainment and presentation rather than true prevalence^44^. Females remain markedly underrepresented in autism neuroimaging research, and this limitation is especially evident in studies of face processing, where many widely cited investigations used predominantly or exclusively male samples. This raises important questions about the generalizability of prior findings and whether sex-related variation in social attention, compensatory strategies, or neural recruitment has been overlooked.

The present study was designed to address these limitations directly. We examined brain activation during the Hariri Emotional Face-Matching Task (EFMT) in a large sample of children and adolescents with and without ASD, deliberately structured to include equal numbers of males and females in both diagnostic groups. This design enables a rigorous test of a central unresolved question in the field: whether autism is associated with reliable differences in neural activation during explicit socioemotional face processing, in a sufficiently powered, developmentally informative, and sex-balanced sample.

The EFMT^45^ is a well-validated fMRI paradigm that robustly engages core and extended face-processing circuitry—including the amygdala, OFA, FFA, and pSTS—by requiring participants to match facial expressions based on emotional content while minimizing linguistic and higher-order cognitive demands^37,46,47^. This makes it particularly well-suited for developmental and clinical populations, as it provides a precise assay of affective face processing while reducing confounds related to language and task complexity. Its widespread use in large-scale, multi-site neuroimaging studies further supports its utility in examining diagnostic differences across heterogeneous developmental samples^48,49^.

In the present study, we leveraged data from Wave 1 of the GENDAAR Project, a multi-site NIH Autism Centers of Excellence (ACE) initiative, to examine neural responses to emotional faces in youth ages 8–18 years. Our primary aim was to test whether ASD diagnosis was associated with altered activation during explicit emotional face processing, particularly within regions implicated in face perception and socioaffective evaluation, including the OFA, FFA, pSTS, and amygdala. Based on prior work, we expected diagnostic differences to be most evident during the processing of emotional facial expressions. However, given the heterogeneity of the existing literature, we treated the magnitude, spatial distribution, and robustness of these effects as key empirical questions. A second aim was to determine whether these effects generalized across sexes in a deliberately sex-balanced developmental sample, thereby providing a more rigorous test of the generalizability of prior findings in autism neuroimaging.

## Methods

### Participants

Data were drawn from Wave 1 of the GENDAAR Project, part of the NIH ACE initiative, which included four MRI acquisition sites: Yale University, Harvard University/Boston Children’s Hospital, University of California, Los Angeles, and University of Washington/Seattle Children’s Research Institute.

As detailed in Table 1, participants (8–18 years, n=265) were recruited into either an autism spectrum disorder (ASD) group (n=128) or a typically developing (TD) comparison group, with equal numbers of both males and females included in each group. Parents provided written informed consent, and participants provided written assent. All procedures were approved by the institutional review boards at each participating site and conducted in accordance with the Declaration of Helsinki.

**Table 1.**
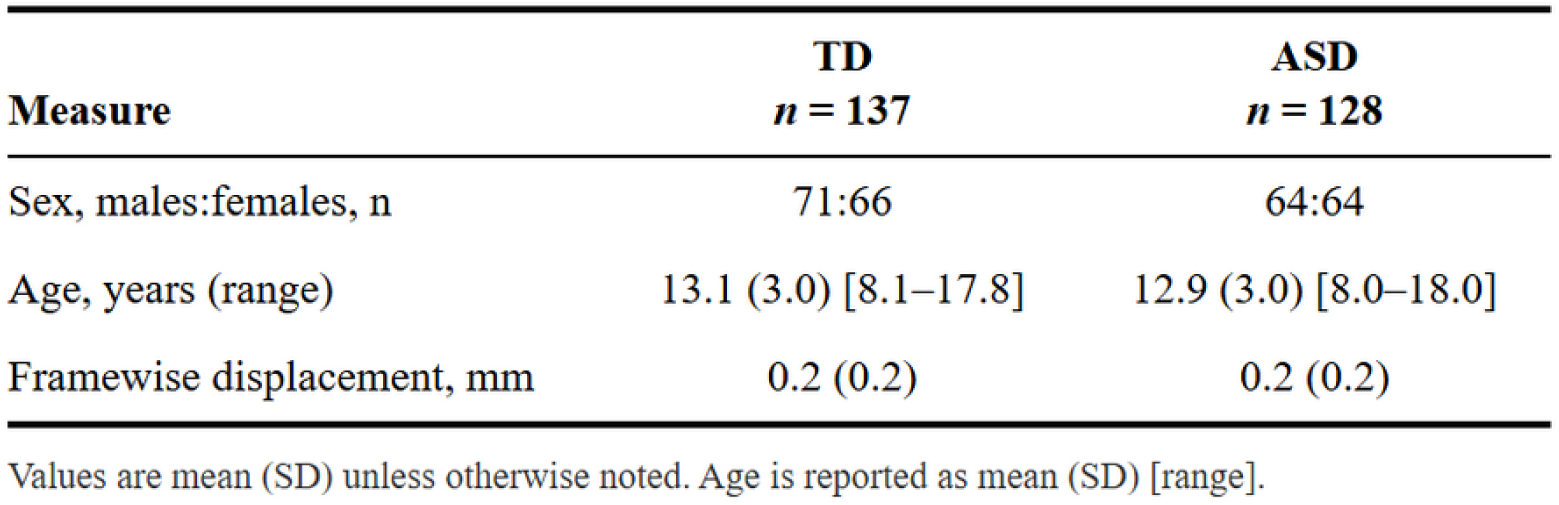
Demographic and clinical characteristics.

| **Measure** | **TD**<br>***n* = 137** | **ASD**<br>***n* = 128** |
| --- | --- | --- |
| Sex, males:females, *n* | 71:66 | 64:64 |
| Age, years (range) | 13.1 (3.0) [8.1–17.8] | 12.9 (3.0) [8.0–18.0] |
| Framewise displacement, mm | 0.2 (0.2) | 0.2 (0.2) |

Values are mean (SD) unless otherwise noted. Age is reported as mean (SD) [range].

Exclusion criteria for all participants included a full-scale IQ ≤ 70 (estimated using the Differential Ability Scales–Second Edition), twin status, neurological or medical conditions that could interfere with imaging (including active tic disorders or recent seizures), pregnancy, presence of metal in the body, and current use of benzodiazepines, barbiturates, or antiepileptic medications. Other medications had to be stable for at least six weeks prior to participation.

ASD diagnoses were confirmed by expert clinicians using the Autism Diagnostic Observation Schedule, Second Edition (ADOS-2; Modules 3 or 4, as appropriate), in conjunction with the Autism Diagnostic Interview–Revised (ADI-R).

### Neuroimaging Data Acquisition

Neuroimaging data were collected across four sites: UCLA, Yale, Harvard, Seattle Children’s, using harmonized acquisition protocols. All sites initially acquired functional MRI data on Siemens 3T Tim Trio scanners; during the study, two sites upgraded to Siemens 3T Prisma Fit systems. To account for potential scanner-related variability, site/scanner was modeled as a nuisance regressor in all group-level analyses, and data acquired before and after scanner upgrades were treated as separate sites.

Whole-brain functional images were acquired with gradient-echo echo-planar imaging sequences harmonized across sites (TR = 2000 ms, TE ≈ 30 ms, flip angle ≈ 90°, voxel size ≈ 3 mm isotropic) in accordance with the standardized ACE Wave 1 protocol (see supplementary methods). Participants viewed stimuli through MRI-compatible video goggles and provided responses via a button box with the left and right index fingers.

### Experimental Paradigm

Participants completed a well-validated emotional face-matching paradigm adapted from Hariri and colleagues^45^. On each trial, a target stimulus was presented above two choice stimuli, and participants were instructed to select the option that matched the target.

In the faces condition, stimuli consisted of fearful and angry facial expressions from the standardized Ekman face set^50^. In the shapes condition, participants matched geometric shapes using the same format, serving as a perceptual control condition. The task included six face blocks and six shape blocks, each with six trials (5 s per trial), yielding 30-s blocks and a total task duration of approximately 6 minutes.

The primary contrast of interest (Faces > Shapes) isolates activity associated with emotional face processing while controlling for low-level perceptual and motor demands.

### Preprocessing and First-Level Analysis

Functional MRI data was preprocessed using fMRIPrep v21.0.2 followed by standard FSL (v6.0.7.22) pipelines, including motion correction, slice-timing correction, spatial smoothing, and denoising^51^. Motion-related artifacts were addressed with motion regressors, scrubbing of high-motion volumes, and ICA-based denoising with ICA-AROMA. Additional preprocessing steps followed best practices for multi-site harmonization within the ACE framework.

At the individual-subject level, task-related neural responses were modeled using a general linear model (GLM) with regressors corresponding to the Faces and Shapes conditions. Contrast of parameter estimate (COPE) images were generated for the Faces > Forms contrast for each participant.

### Pre-Defined Region-of-Interest Definition

Analyses focused on *a priori* regions implicated in face processing, using a hybrid region-of-interest (ROI) approach. Functional masks for the right FFA, right OFA, and right pSTS were derived from Neurosynth meta-analytic maps with the search terms “face processing” and “posterior temporal sulcus.” The left and right amygdala were defined anatomically using the Harvard–Oxford atlas in FSL. This process yielded a total of 5 distinct regions of interest. Given that face processing is generally right latera lized^8,9,10^, the analyses focused on voxels from the right hemisphere for each ROI. Conversely, amygdala activation during face processing has been shown to be bilaterial^20,45,52^, so voxels from both hemispheres were combined for the amygdala ROI.

### Group-Level Statistical Analysis

Group-level analyses were conducted using mixed-effects models in FSL, with supplementary analyses performed in R (v4.5.0). The Faces > Shapes contrast was examined across diagnostic groups (ASD vs. TD) and sex groups (male vs. female). Although both the identity and match conditions involved faces, they were analyzed separately due to preliminary analyses showing differential activation patterns across the key ROIs. Furthermore, matching and identifying faces may require distinct pathways; matching shows greater bilateral amygdala activation while identifying engages the prefrontal circuits^45,53,54^. Mean activation was computed within each subgroup, and planned comparisons included within-sex diagnostic contrasts (e.g., TD females > ASD females), within-diagnosis sex contrasts (e.g., ASD females > ASD males), and their interaction.

To ensure robust statistical inference, nonparametric permutation testing was performed using FSL’s randomise with threshold-free cluster enhancement (TFCE), 10,000 permutations, and family-wise error correction (p < .05), following recommendations by Eklund and colleagues^55^.

All models included age (grand-mean centered), GCA (grand-mean centered), and site (dummy coded) as covariates. As shown in Table 2, the groups differed in on GCA. As such, GCA was also included as a covariate. Sex and diagnosis were modeled as fixed effects.

**Table 2.** Clinical measures by sex and diagnosis.

| **Measure** | **TD**<br>**Female**<br>***n* = 66** | **TD**<br>**Male**<br>***n* = 71** | **ASD**<br>**Female**<br>***n* = 64** | **ASD**<br>**Male**<br>***n* = 64** |
| --- | --- | --- | --- | --- |
| IQ (GCA) | 112.6<br>(14.2) | 115.1<br>(15.4) | 101.9<br>(21.9) | 100.9<br>(18.8) |
| SRS total raw score | 17.4 (11.9) | 17.5 (15.2) | 93.3 (29.0) | 91.3 (25.2) |
| ADOS comparison score | — | — | 6.2 (2.1) | 6.8 (2.2) |
| ADI-R Social total | — | — | 18.6 (6.4) | 19.0 (5.3) |
| ADI-R Communication<br>total | — | — | 15.3 (4.8) | 16.5 (4.0) |
| ADI-R Behavioral total | — | — | 5.6 (2.8) | 6.4 (2.6) |

Values are mean (SD). GCA = General Conceptual Ability; SRS = Social Responsiveness Scale; ADOS = Autism Diagnostic Observation Schedule; ADI-R = Autism Diagnostic Interview–Revised.

## Results

As illustrated in Figure 1, the ASD group showed less activation relative to the TD group for the Faces>Forms contrast. In the right fusiform face area, there were significant activation results due to group (p=0.02) and sex (p=0.007) individually, as well as a group by condition interaction (p=0.01). In the group by condition interaction, there was significantly lower activation in ASD males than in ASD females, TD females, and TD males. The activation difference between ASD males and TD males was significant. Collapsing across conditions, there is a significant group by sex interaction within the ASD male and female groups.

**Figure 1:**
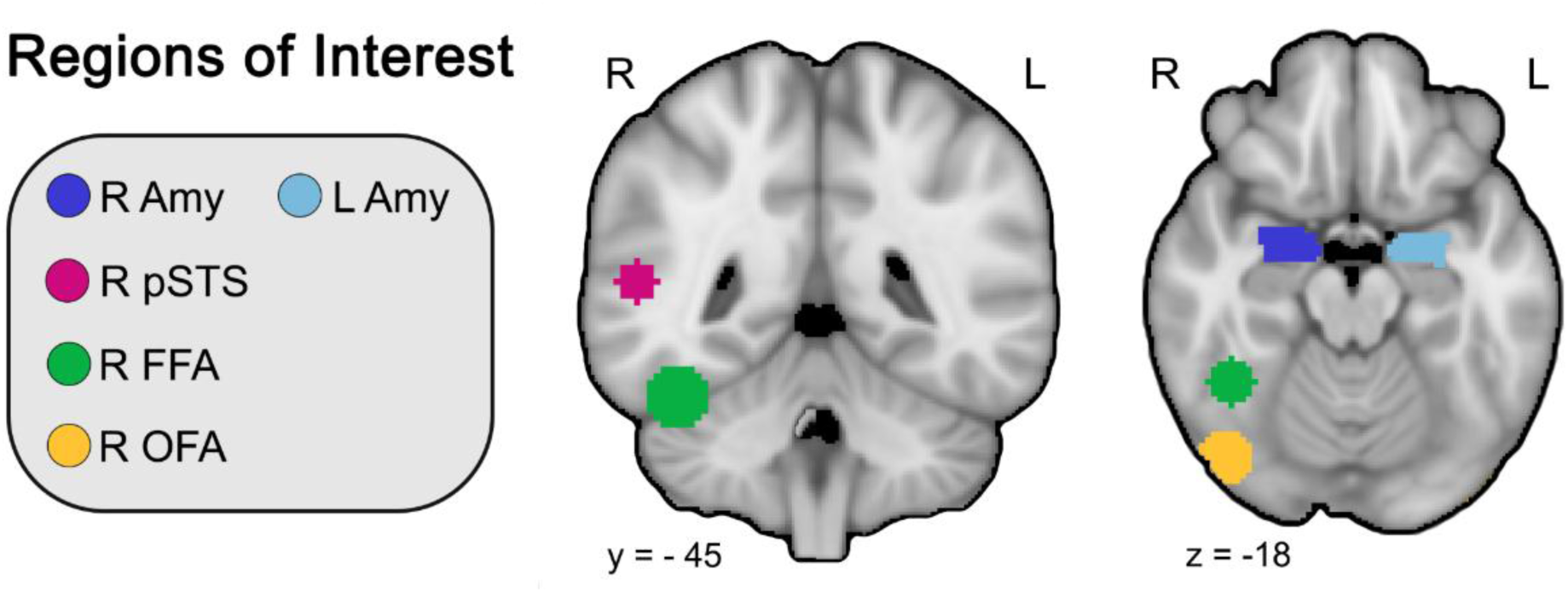
Anatomical masks of selected ROI

**Figure 2:**
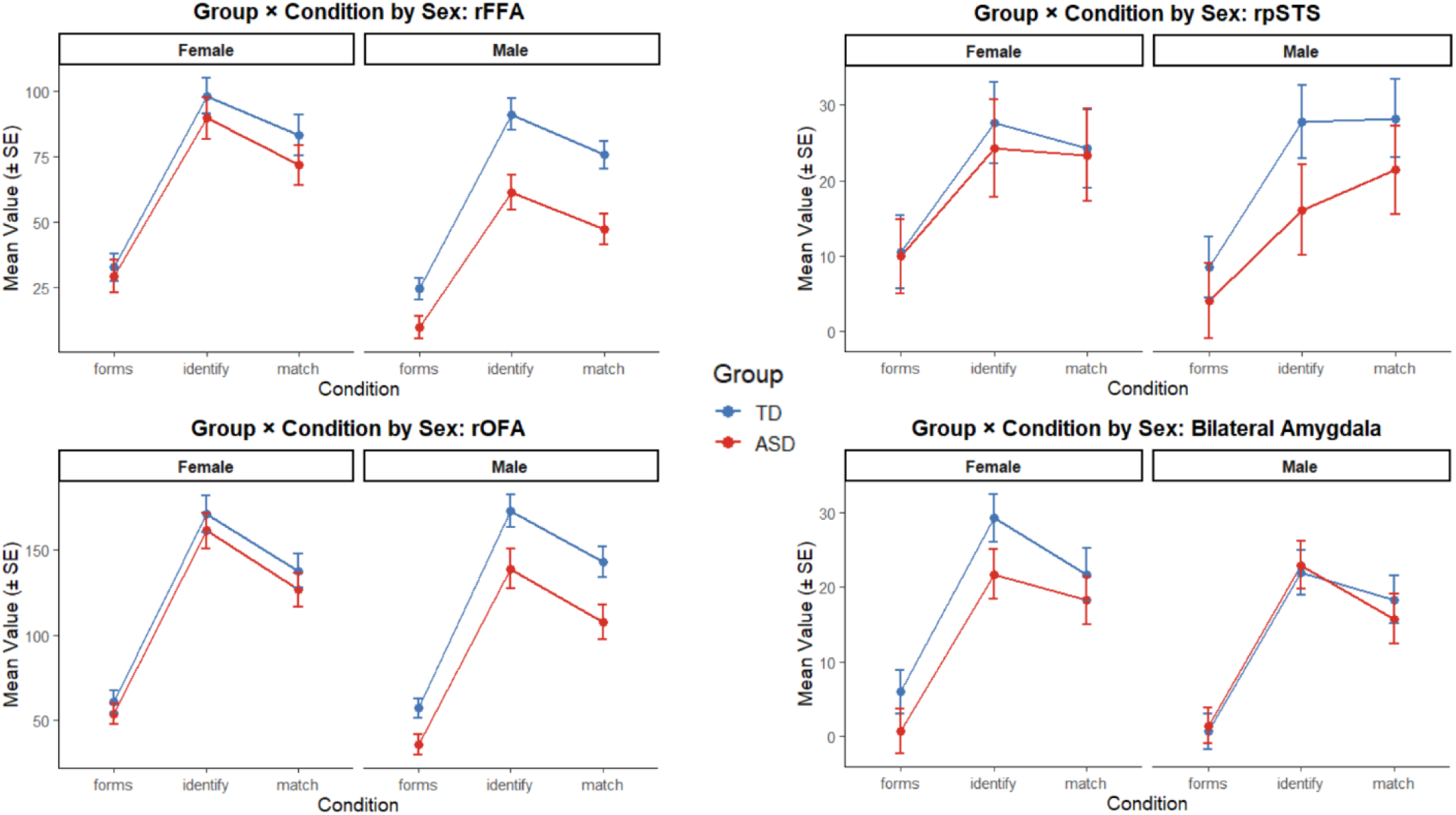
Group by condition interaction presented by sex for the rFFA, rOFA, rpSTS, and bilateral amygdala

## Discussion

The present study examined neural responses during explicit emotional face processing in a large, developmentally informative, and sex-balanced sample of youth with and without ASD. Consistent with prior work, we observed reduced activation in ASD relative to typically developing participants within components of the canonical face-processing network, particularly the right fusiform face area. However, these effects were neither global nor uniform across groups. Instead, the most robust differences emerged among autistic males, who showed significantly reduced activation relative to autistic females and typically developing participants. These findings suggest that atypical neural responses to emotional faces in autism may be more selective, heterogeneous, and sex-dependent than early models of a generalized “social brain deficit” proposed.

Importantly, the overall pattern of findings both converges with and refines prior neuroimaging research on face processing in autism. Early influential fMRI studies frequently reported reduced fusiform activation during face perception tasks and interpreted these findings as evidence of disrupted social specialization within ventral visual cortex^23,24^. Over time, however, the literature became increasingly mixed, with some studies observing diminished activation^23–25^, others finding relatively intact responses under controlled attentional conditions^27,38,41^, and still others reporting evidence for compensatory recruitment or alternative neural strategies^24,31^. The present findings help reconcile some of this inconsistency. In a relatively large and rigorously characterized developmental sample, we did not observe pervasive reductions across the broader face-processing network. Rather, effects were localized and moderated by sex, suggesting that prior inconsistencies may partly reflect differences in sample composition and insufficient representation of females.

The findings within the fusiform face area are particularly noteworthy given the longstanding centrality of this region in autism neuroimaging models. The right FFA is critically involved in the perceptual encoding of faces and is typically strongly engaged during emotional face matching tasks^8,12,37,46^. In the current study, autistic males showed significantly lower activation in this region relative to both autistic females and typically developing males, whereas autistic females showed substantially less divergence from comparison participants. This pattern raises the possibility that at least some prior reports of reduced fusiform activation in autism may have been disproportionately driven by predominantly male samples, which historically characterized much of the literature. Historically, autism research has overrepresented males in study populations, at times with male:female ratios of 4:1 up to 8:1, leading to generalization of findings that may be male-specific^56^.

These findings contribute to a growing body of evidence suggesting that sex may meaningfully shape the neural phenotype of autism. Behavioral and clinical studies increasingly indicate that autistic females often differ from autistic males in patterns of social attention^57,58^, compensatory behavior^59,60^, symptom presentation ^61,62^, and ascertainment^63,64^. Neuroimaging work examining these questions remains limited, however, due largely to the persistent underrepresentation of females in autism research. By deliberately balancing the sample across diagnosis and sex, the present study provides a more rigorous test of whether neural differences associated with autism generalize across sexes. The results suggest that they may not fully do so. Instead, the neural correlates of emotional face processing in autism appear at least partly moderated by sex, underscoring the importance of moving beyond male-dominant models of autism neurobiology.

At the same time, the present findings also argue against overly simplistic deficit-based accounts of social perception in autism. Despite statistically significant group differences, autistic participants as a whole robustly engaged core face-processing circuitry during the task. This is an important point. The EFMT is an explicit socioemotional matching paradigm that strongly recruits face-sensitive and salience-related systems in most individuals^45,46,65^. The observation that many autistic youth showed substantial activation within these networks suggests that the neural systems supporting face perception are not absent or fundamentally nonfunctional in autism. Rather, differences may emerge in the efficiency, modulation, developmental tuning, or contextual deployment of these systems. Such an interpretation is more consistent with contemporary dimensional and developmental models of autism than with earlier notions of a unitary social perceptual deficit^66,67^. Neural activation may exist on a spectrum that overlaps significantly with typically developing individuals, and placement may be strongly associated with sex^68^.

The relatively selective nature of the observed effects may also reflect important features of the task itself. The EFMT minimizes linguistic and higher-order cognitive demands while explicitly directing participants’ attention toward emotional facial expressions. Prior work suggests that attentional engagement and gaze allocation can substantially influence fusiform and amygdala responses in autism^27,69–71^. Thus, paradigms that require explicit attention to faces may reduce some group differences relative to passive viewing tasks or more naturalistic social paradigms. The present findings therefore highlight the importance of considering task demands carefully when interpreting inconsistencies across the literature.

Several limitations should be considered. First, although the sample was substantially larger and more sex-balanced than many prior neuroimaging studies of autism, the study remains cross-sectional and cannot directly characterize developmental trajectories of face-processing circuitry over time. Similar phenotypic outcomes may result from drastically different developmental pathways. Longitudinal work will be important for determining whether the observed sex-related differences reflect distinct developmental pathways, compensatory adaptations, or differences in social experience across development. Specifically in females, longitudinal studies can help determine if atypical recruitment of frontoparietal control networks^72^ works to train the brain to respond to social stimuli appropriately^73^. Social experiences enforce traditional gender roles and social interaction standards that subject females to increased conformational pressure, that may modulate face processing activation over time^57,74^. Second, although the EFMT robustly probes emotional face processing, it does not directly measure gaze behavior, visual attention, or social motivation during task performance^5,27,75,76^. Future studies integrating eye tracking and naturalistic paradigms may help clarify the mechanisms underlying individual differences in neural activation. Third, the current analyses focused on regional activation differences within predefined ROIs. Functional connectivity and multivariate approaches may reveal additional differences in network organization or representational structure not captured by univariate activation analyses alone. For example, the recruitment of additional processing pathways and accessory regions of the brain.

Despite these limitations, the present study addresses several longstanding gaps in the autism neuroimaging literature. By leveraging data from the GENDAAR Project, we examined emotional face processing in one of the larger developmental fMRI samples to include balanced representation of autistic males and females. The findings suggest that neural differences during socioemotional face processing in autism are present but more nuanced than early models proposed, with evidence for selective alterations centered particularly within right fusiform cortex and moderated by sex. More broadly, the results underscore the importance of adequately powered, developmentally informed, and sex-balanced neuroimaging studies for refining models of autism neurobiology.

Ultimately, understanding variability in the neural systems supporting social perception may be critical for advancing more individualized and mechanistic accounts of autism. Rather than asking whether autistic individuals do or do not recruit “typical” face-processing circuitry, future work may benefit from focusing on how developmental trajectories, sex-related variation, attentional processes, and broader neurobiological heterogeneity shape the organization of social brain systems across individuals on the autism spectrum.

